# Real-time luminescence assay for cytoplasmic cargo delivery of extracellular vesicles

**DOI:** 10.1101/2020.10.16.341974

**Authors:** Masaharu Somiya, Shun’ichi Kuroda

**Author notes:** Corresponding author: Prof. Masaharu Somiya, Ph.D., Department of Biomolecular Science and Reaction, The Institute of Scientific and Industrial Research, Osaka University, 8-1 Mihogaoka, Ibaraki, Osaka 567-0047, Japan.

## Abstract

Extracellular vesicles (EVs) have been considered to deliver biological cargos between cells and mediate intercellular communication. However, the mechanisms that underlie the biological process of EV uptake and cytoplasmic cargo release in recipient cells are largely unknown. Quantitative and real-time assays for assessment of the cargo delivery efficiency inside recipient cells have not been feasible. In this study, we developed an EV cargo delivery (EVCD) assay using a split luciferase called the NanoBiT system. Recipient cells expressing LgBiT, a large subunit of luciferase, emit luminescence when the EV cargo proteins fused with a small luminescence tag (HiBiT tag) that can complement LgBiT are delivered to the cytoplasm of recipient cells. Using the EVCD assay, the cargo delivery efficiency of EVs could be quantitatively measured in real time. This assay was highly sensitive in detecting a single event of cargo delivery per cell. We found that modification of EVs with a virus-derived fusogenic protein significantly enhanced the cytoplasmic cargo delivery; however, in the absence of a fusogenic protein, the cargo delivery efficiency of EVs was below the threshold of the assay. The EVCD assay could assess the effect of entry inhibitors on EV cargo delivery. Furthermore, using a luminescence microscope, the cytoplasmic cargo delivery of EVs was directly visualized in living cells. This assay could reveal the biological mechanism of the cargo delivery processes of EVs.

## Introduction

Extracellular vesicles (EVs), membranous nanoparticles secreted by living cells, are thought to be involved in intercellular communication in various species from microorganisms to vertebrate ^1,2^. Since EVs contain cargo molecules such as RNAs and proteins in their luminal space, they may deliver the cargo molecules into recipient cells and regulate biological functions in the recipient cells. Numerous studies have shown that the treatment of recipient cells with EVs containing specific cargos (especially microRNAs or proteins) results in phenotypic changes in the recipient cells. Owing to the delivery capability of biomolecules, EVs have been studied as a promising drug delivery system for therapeutic proteins or RNAs ^3,4^.

However, the cargo delivery mechanism of EVs, especially the process of cytoplasmic cargo release, remains largely unknown ^5^. Mechanistically, EVs are mainly endocytosed by recipient cells, fuse with the endosomal/lysosomal membrane, and release their cargo into the cytoplasm ^5,6^. Although few studies have shown that EVs are capable of fusing with the cellular membrane of recipient cells ^7,8^, direct evidence indicating the cytoplasmic cargo delivery of EVs has not been demonstrated.

We discussed the possibility of “EV cargo transfer hypothesis” in our previous review and concluded that cargo delivery by EVs might not be a frequent event as generally accepted ^9^. Several studies have suggested that EV-mediated cargo delivery is a rare event. When the recipient cells are treated with EVs *in vitro*, only 0.1% to 5.0% of the cell population exhibit the functional readout of cargo delivery, although the efficacy depends on the experimental system ^10–12^.

To decipher the mechanism and physiological relevance of EV cargo delivery, a feasible and reliable assay to measure cargo delivery in real time is necessary. Conventionally, cargo delivery of EVs is evaluated by the phenotypic change in the recipient cells, although these methods are often interfered with the experimental artifacts that can be induced by contaminants in the EV fraction ^13^. Another approach for the assessment of EV cargo delivery involves use of reporter assays for measuring functional miRNA activity in recipient cells ^14^. This assay is based on the assumption that EV-mediated delivery of miRNA leads to a change in reporter gene expression in recipient cells. However, this assay could not demonstrate direct evidence of cargo transfer by EVs because of several confounding factors ^9^.

In this study, we developed a quantitative and real-time luminescence assay to measure cargo protein delivery by EVs in recipient cells. The key feature of this EV cargo delivery (EVCD) assay is the luciferase complementation assay using *Oplophorus gracilirostris*-derived highly bright luciferase (NanoLuc) ^15,16^. A small fragment of NanoLuc (HiBiT tag) was fused to EV cargo proteins, while the large subunit of NanoLuc (LgBiT) was expressed in recipient cells. When the HiBiT-tagged cargo proteins are delivered to the cytoplasm of recipient cells, luciferase fragments complement and emit luminescence signals (Fig. 1A). Since the complemented NanoLuc is much brighter than conventional luciferases such as firefly or *Renilla* luciferases, NanoLuc-based assays are sensitive enough to detect the rare event of EV cargo delivery. Furthermore, this assay enabled us to measure the kinetics of cargo protein delivery by EVs and to visualize the cytoplasmic cargo delivery by EVs in real time.

**Fig. 1.**
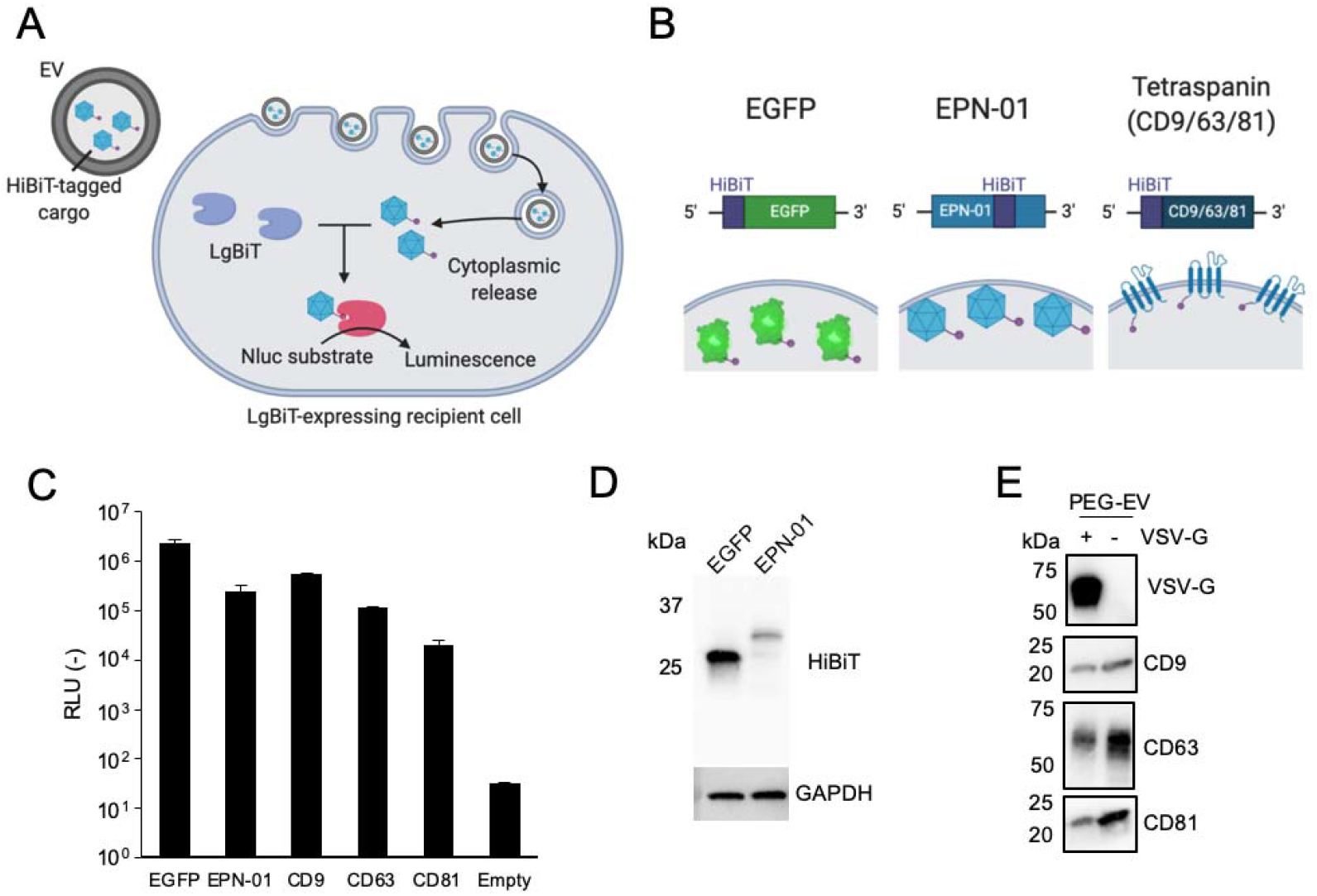
Summary of the EVCD assay and characterization of HiBiT-tagged EV cargo proteins. (A) Schematic representation of the EVCD assay. EV containing HiBiT-tagged cargo is internalized by LgBiT-expressing recipient cells, followed by cytoplasmic release of the cargo. Spontaneous complementation of HiBiT-tagged protein cargo with LgBiT leads to the elicitation of luminescence. (B) Schematic representation of HiBiT-tagged proteins. Upper panels show the structure of expression plasmids. Lower panels show the protein localization inside EVs. (C) Expression levels of HiBiT-tagged EV cargo proteins in donor HEK293T cells. N=3, mean ± SD. (D) Detection of HiBiT-tagged EGFP and EPN-01 in cell lysate of transfected HEK293T cells using LgBiT as a probe. GAPDH was used as a loading control. (E) Detection of VSV-G and EV marker proteins in purified EV fraction. Culture supernatant was concentrated by PEG precipitation and subjected to western blotting

## Results and Discussion

### Characterization of HiBiT-tagged EV cargo proteins

The EVCD assay (Fig. 1A) is based on the complementation of HiBiT and LgBiT in the cytoplasm. When the EVs containing HiBiT-tagged cargo are delivered to the cytoplasm of LgBiT-expressing recipient cells, emitted luminescence can be detected. To establish the EVCD assay, we first attempted to tag the EV cargos with HiBiT (Fig. 1B). Three types of protein EV cargos including EGFP, a self-assembling protein EPN-01 ^17^, and tetraspanins were used. The first cargo EGFP was tagged at the N-terminal with HiBiT and overexpressed in the donor cells. Cytoplasmic EGFPs may be passively loaded into EVs. The second cargo EPN-01 was a nanocage-forming protein that was designed *de novo* and secreted from cells via the ESCRT pathway with EVs ^17^. The original Myc-tag of this cargo was replaced with an HiBiT tag. Tetraspanins, typical EV marker proteins embedded in the EV membrane, were also tagged with HiBiT at their N-termini.

All HiBiT-tagged proteins were expressed in HEK293T cells, and expression levels were measured by mixing cell lysates with LgBiT and NanoLuc substrates (Fig. 1C). EGFP and EPN-01showed higher expression levels compared to tetraspanins., Expression of HiBiT-tagged EGFP and EPN-01 was confirmed by western blotting using LgBiT as a probe protein (Fig. 1D). These results confirmed that EGFP and EPN-01 had feasibility in the EVCD assay.

We assessed whether HiBiT-tagged cargo proteins were encapsulated in EVs by immunoprecipitation (Fig. S1). EVs in the culture supernatant were immunoprecipitated using antibodies targeting CD81, a typical EV marker (Fig. S1A), and vesicular stomatitis virus glycoprotein (VSV-G), a fusogenic viral membrane protein that was incorporated into EVs (Fig. S1B). When the supernatant of the cargo protein-expressing cells was immunoprecipitated using anti-CD81 antibodies, cargo proteins were precipitated, indicating that cargo proteins were encapsulated inside EVs (Fig. S1C and S1D). Furthermore, anti-VSV-G antibody could enrich HiBiT-tagged proteins co-expressed with VSV-G, suggesting the incorporation of VSV-G in the EV membrane ^17^ and encapsulation of HiBiT-tagged cargo proteins (Fig. S1C and S1E). Immunoprecipitation was strongly abrogated by detergent treatment (Fig. S1F), indicating that the HiBiT-tagged cargo protein was encapsulated in VSV-G^+^ and/or CD81^+^ membrane vesicles.

Generally, the amount of EVs in the supernatant is low; therefore, a concentration process is necessary to acquire a sufficient amount of EVs for the assay. In this study, EVs containing HiBiT-tagged proteins were concentrated by poly(ethylene glycol) (PEG) precipitation, which is a feasible concentration method for small-scale purification ^18^. As shown in Table 1, HiBiT-tagged EPN-01 was enriched more than 10-fold by PEG precipitation, whereas HiBiT-tagged EGFP was not enriched. This result indicated that a large fraction of EPN-01 in the supernatant was encapsulated within EVs, while the majority of EGFPs were not encapsulated in EVs. Concentrated EV fraction contained VSV-G and EV marker proteins (CD9, CD63, and CD81), suggested that PEG precipitation successfully concentrated the VSV-G-displaying EVs (Fig. 1E).

**Table 1.** PEG precipitation of EVs encapsulating HiBiT-tagged protein cargo (N = 3, mean ± SD)

| Transfection |  | HiBiT enrichment (-)<sup>1</sup> | HiBiT yield (%) |
| --- | --- | --- | --- |
| EPN-01 | - | 12.4 ± 3.4 | 27.6 ± 7.6 |
| | + VSV-G | 12.4 ± 1.4 | 27.5 ± 3.1 |
| EGFP | - | 0.3 ± 0.1 | 3.3 ± 2.5 |
| | + VSV-G | 0.5 ± 0.3 | 6.0 ± 5.4 |

<sup>1</sup> HiBiT enrichment factor was calculated using the amounts of HiBiT before and after the PEG precipitation.

Notably, substantial amounts of non-encapsulated HiBiT-tagged proteins (both EPN-01 and EGFP) were present in the resultant EV fraction that could interfere with the EVCD assay. Therefore, in the subsequent EVCD assay, it is mandatory to use a DrkBiT peptide that complements and inactivates the luciferase activity of LgBiT to competitively block the non-encapsulated HiBiT-tagged proteins (see below).

### Real-time EVCD assay

We first estimated the sensitivity of the EVCD assay using a synthetic HiBiT peptide. After lysis of approximately 1.0 × 10^5^ LgBiT-expressing HEK293T cells, an HiBiT peptide and a NanoLuc substrate were added to the lysate and luminescence signal was measured (Fig. 2A). Approximately 0.1 fmol of the HiBiT peptide was detected, suggesting that the assay was capable of measuring remarkably less amounts of cytoplasmic cargo in the recipient cells.

**Fig. 2.**
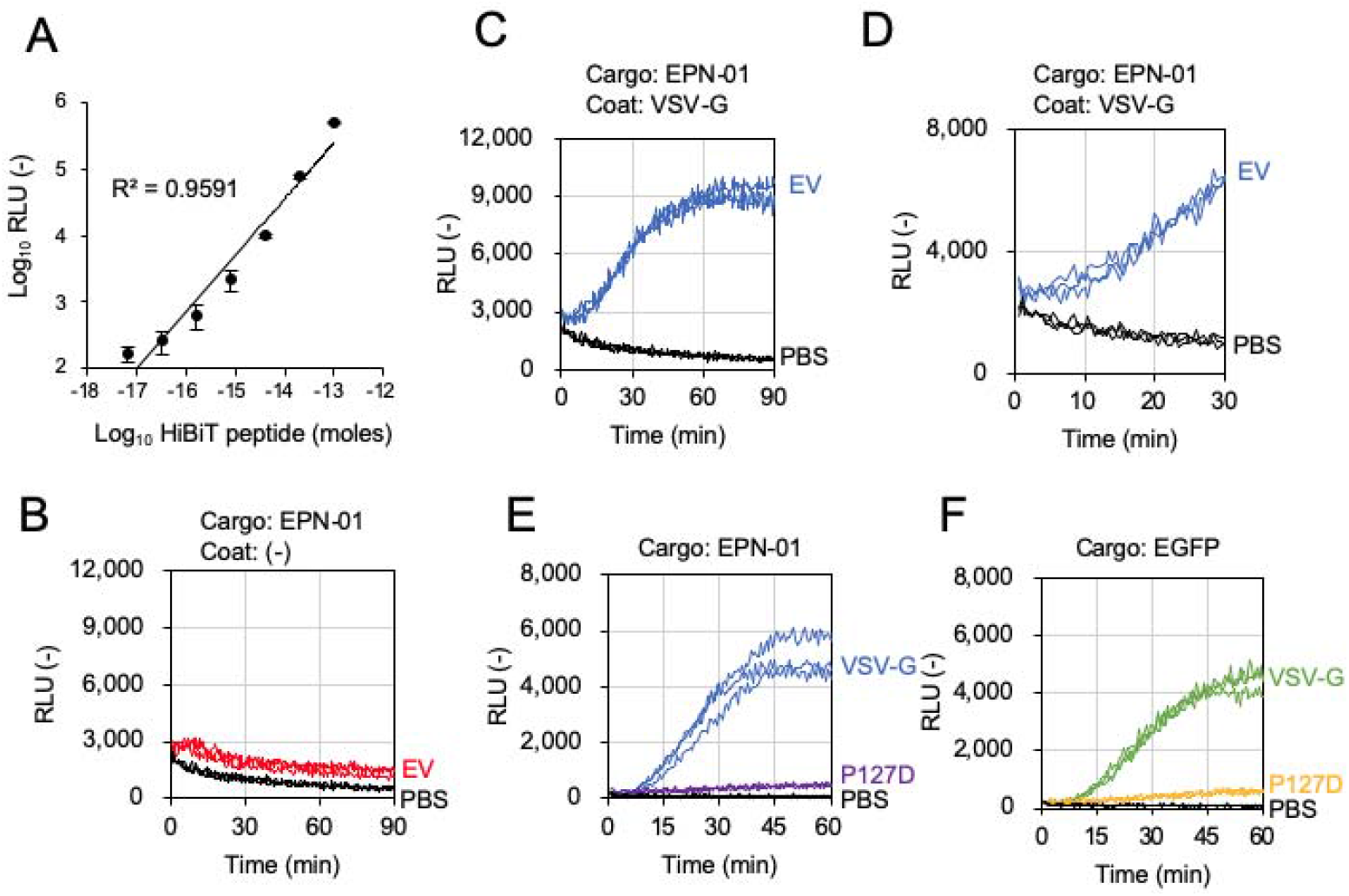
EV cargo delivery (EVCD) assay. (A) Quantitative curve of the HiBiT peptide in cell lysate of LgBiT-expressing HEK293T. N=3, mean ± SD. (B) EVCD assay using EPN-01-containing EVs without fusogenic protein. (C) EVCD assay using EPN-01-containing EVs with fusogenic protein VSV-G. (D) Enlargement of (C) from 0 to 30 min. (E) EVCD assay using EPN-01-containing EVs with either VSV-G or VSV-G(P127D). (F) EVCD assay using EGFP-containing EVs with either VSV-G or VSV-G(P127D). PBS was used as a negative control. All kinetics data represent information obtained from experiments conducted in triplicate.

Next, we measured the cargo delivery kinetics of EVs containing either HiBiT-tagged EGFP or EPN-01. The first observation of EPN-01-containing EVs in the EVCD assay demonstrated no luminescence signal within 90 min (Fig. 2B). For controls, we used the EVs displaying VSV-G proteins, which confer EVs with fusogenic activity that facilitates the cargo delivery of EVs by membrane fusion between the EV and cellular membranes ^17,19^. Evidently, VSV-G-displaying EVs induced a gradual increase in the luminescence signal (Fig. 2C), suggesting that the HiBiT-tagged EPN-01 was delivered to the cytoplasm and the presence of membrane fusion proteins such as VSV-G was indispensable for achieving substantial cargo delivery. The luminescence signal was observed as soon as 20 min after the addition of EVs (Fig. 2D), suggesting that VSV-G could induce prompt fusion and release of EPN-01 cargo into the cytoplasm. This result was consistent with that of previous studies showing that the internalization and fusion of VSV was a rapid process, within 3 min in HeLa cells ^20^ and 20 min in BHK cells ^21^.

As described above, the concentrated EV fraction contains a substantial amount of HiBiT-tagged proteins outside of EVs. Moreover, LgBiT may be leaked from the recipient cells into the medium ^22^. These components significantly affect the sensitivity and accuracy of the EVCD assay. As shown in Fig. S2, in the absence of DrkBiT, a sudden increase in the luminescence signal was observed immediately after the addition of EVs (Fig. S2A). However, in the presence of 1 μM DrkBiT in the buffer, luminescence signal emitted by EV-mediated cargo delivery was distinguishable from non-specific luminescence signal (Fig. S2B), indicating that nonspecific complementation of LgBiT and HiBiT outside the recipient cells interfered with the assay. Therefore, it is mandatory to use a DrkBiT peptide in the EVCD assay to avoid non-specific background signals.

To validate the EVCD assay and exclude an experimental artifact, we used mutant VSV-G(P127D) that is incapable of fusing with the host cell membrane ^23^. Using both EGFP and EPN-01 as cargos, VSV-G(P127D) decreased the cargo delivery efficacy of EVs compared to the parental VSV-G (Fig. 2D and 2E), which was consistent with the findings of a previous report demonstrating that the fusogenic activity of VSV-G was indispensable for cytoplasmic delivery of the EV cargo ^17,24^. These results support that the EVCD assay can elucidate the fusion and cytoplasmic cargo release of EVs in recipient cells.

### Evaluation of EV entry inhibitors using the EVCD assay

We evaluated the effect of compounds that are known to inhibit endocytosis and membrane fusion by using the EVCD assay with EPN-01-containing EVs modified with VSV-G. Chlorpromazine ^25^, Dynasore ^26^, EIPA ^27^, and Pitstop 2 ^28^ have been known to inhibit the endocytosis of EVs, and all these compounds could significantly decrease the cargo delivery of EVs (Fig. 3A-3D). Furthermore, chloroquine ^29^ and bafilomycin A1 ^30^, both known to inhibit low pH-dependent fusion activity of VSV-G, abolished the cargo delivery of EVs in a dose-dependent manner (Fig. 3E and 3F). These results confirmed that the EVCD assay could evaluate the cargo delivery efficiency of EVs and the effect of inhibitors.

**Fig. 3.**
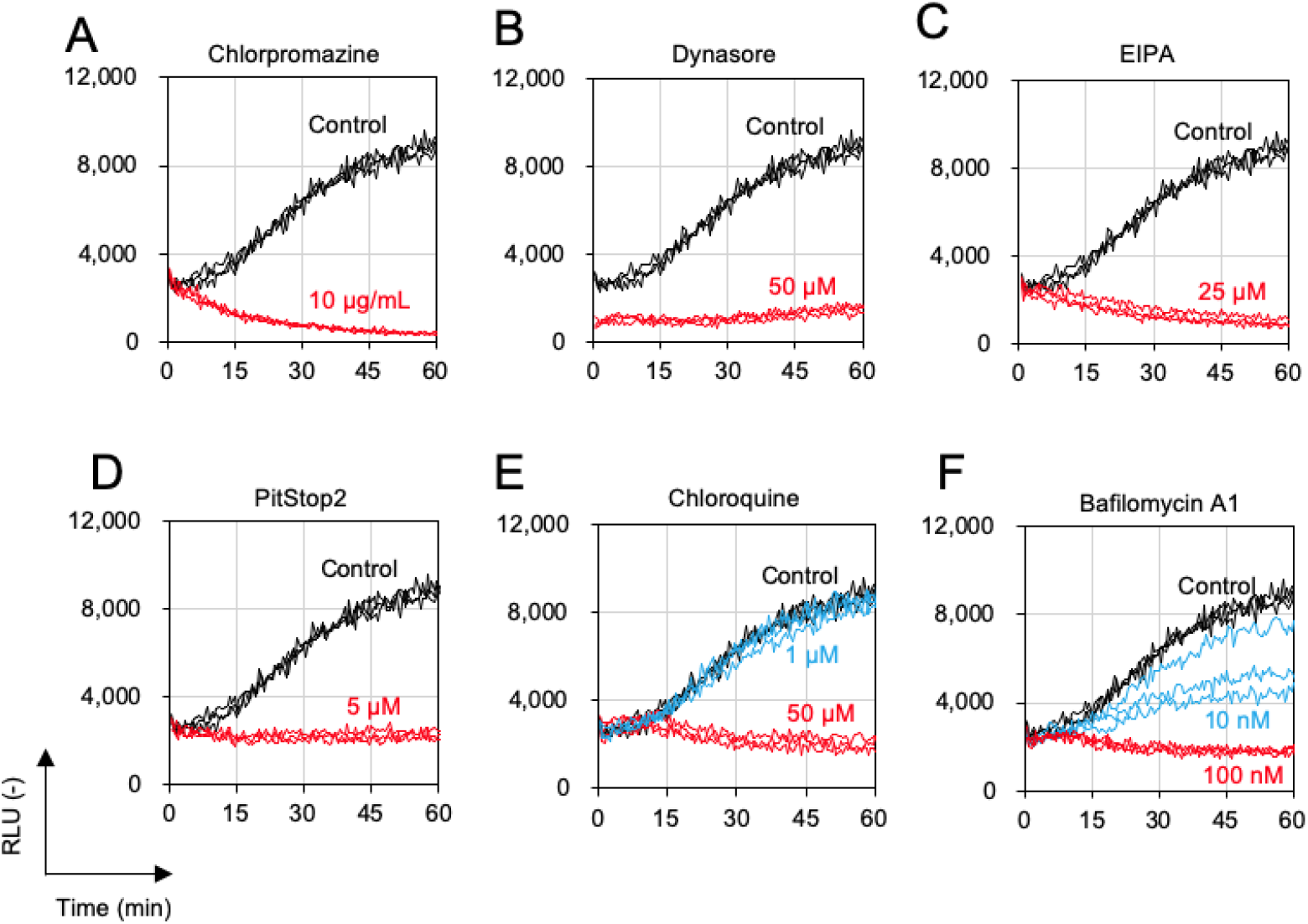
The inhibitory effect of compounds on cargo delivery by EVs. Endocytosis inhibitors (chlorpromazine [A], Dynasore [B], EIPA [C], and Pitstop 2 [D]) and membrane fusion inhibitors (chloroquine [E] and bafilomycin A1[F]) were analyzed in the EVCD assay using EPN-01-containing EVs modified with VSV-G. All kinetics data represent information obtained from experiments conducted in triplicate.

### The use of the EVCD assay to decipher the endosomal escape efficiency of EVs

It has been reported that endosome-destabilizing reagents such as chloroquine and UNC10217832A can enhance the cargo delivery of EVs ^31^. To confirm the effect of the endosomolytic reagent on the cargo delivery efficiency of EVs without VSV-G modification, we evaluated whether chloroquine could enhance the cargo delivery of EVs using the EVCD assay. Unexpectedly, chloroquine did not enhance the cargo EPN-01 delivery by EVs (Fig. 4A) for 90 min. Conversely, cargo delivery of VSV-G-modified EVs was significantly reduced by chloroquine (Fig. 3E and 4B), suggesting that chloroquine increased the pH within endosomes/lysosomes in recipient cells and inhibited membrane fusion by VSV-G.

**Fig. 4.**
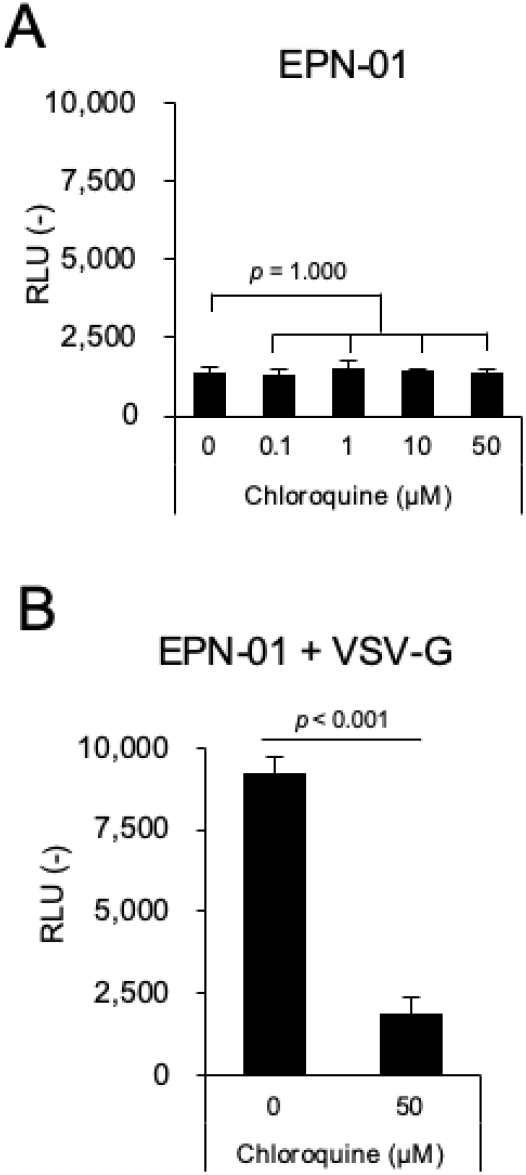
Cargo delivery efficiency of EPN-01-containing EVs in the presence of chloroquine. (A) EVs without VSV-G and (B) EVs with VSV-G. Luminescence signal after 90 min of EV treatment was represented. N=3, mean ± SD. Statistical analysis was performed using one-way ANOVA followed by post hoc Dunnett’s test (A) and the Student’s *t*-test (B).

### Real-time imaging of cytoplasmic cargo delivery by EVs

Imaging of the real-time cytoplasmic delivery of cargo molecules in recipient cells is of prime importance, as the localization and timing of cargo delivery of EVs is largely unknown. The EVCD assay described above can analyze a considerable segment of the event of cargo delivery in a cell population with high sensitivity. Therefore, we attempted to observe the luminescence signal emitted by cargo delivery at the single-cell level using a luminescence microscope. We succeeded in capturing the cytoplasmic cargo release of VSV-G-containing EVs in LgBiT-expressing HEK293T cells (Fig. 5A to 5C and Supplementary Video). As shown in Fig. 5B and 5C, luminescence dots were observed within recipient cells over time, suggesting that EPN-01 nanocages were anchored to the cytoplasmic leaflet of the endo/lysosomal membrane as an intact nanocage (Fig. 5D) because of the N-terminal myristoyl group of EPN-01 that could anchor the membrane organelle ^17^.

**Fig. 5.**
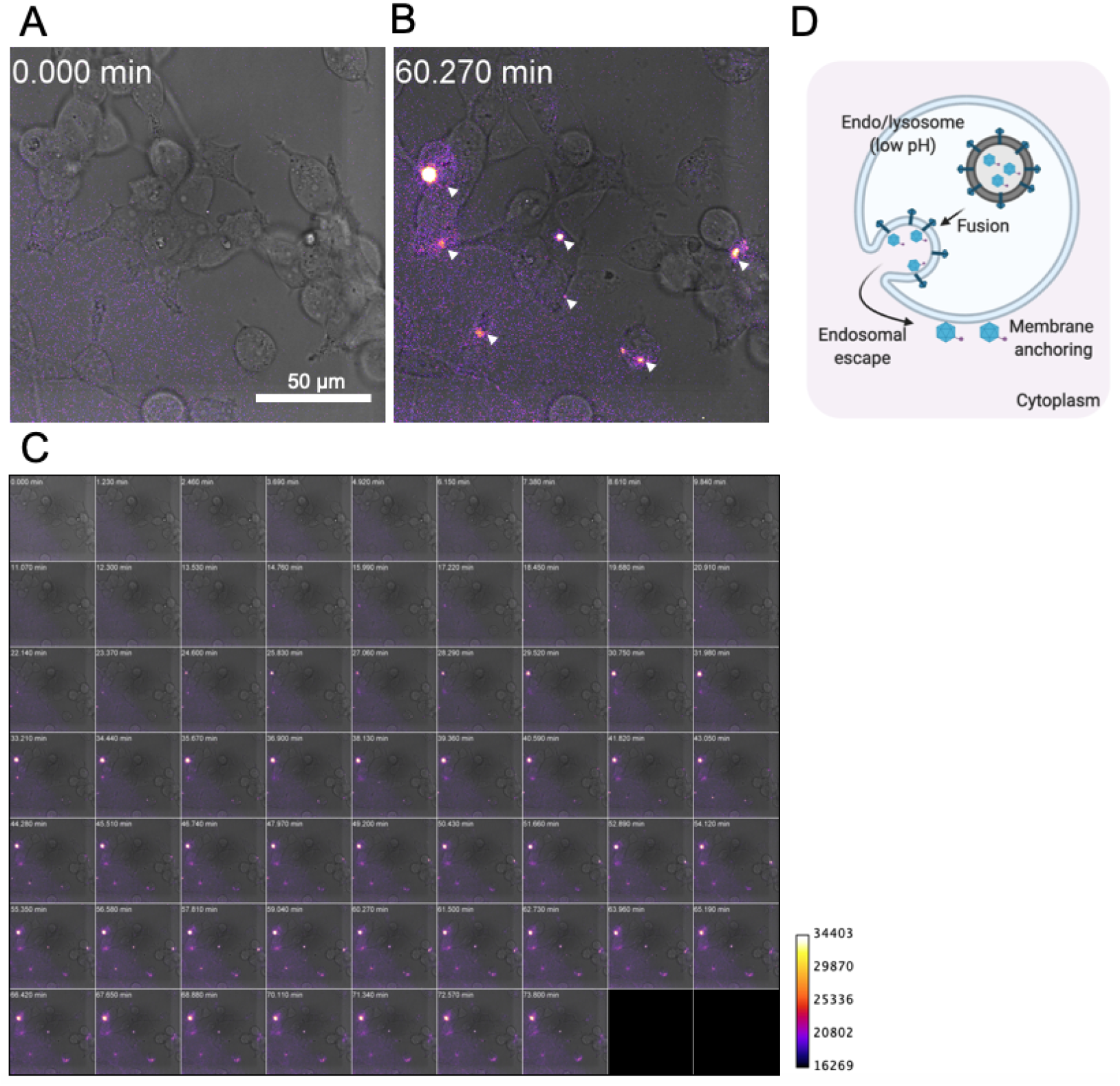
Live cell imaging of cargo delivery of EVs with VSV-G. Luminescence images of recipient HEK293T cells before (A) and after 60 min (B) of treatment with EPN-01-containing EVs modified with VSV-G. Arrowheads indicate complemented NanoLuc-derived luminescence signals within cells. (C) Series of luminescence images of cells treated with EPN-01-containing EVs modified with VSV-G from 0 to 73.8 min (see Supplementary Video). EVs were added at 0 min. (D) Expected intracellular localization of EPN-01 after the fusion of EVs and cytoplasmic release.

## Conclusions

In this study, we developed a novel assay to measure the real-time cargo delivery efficiency of EVs in living recipient cells. Previously, the NanoBiT technology has been used to evaluate viral entry ^22,32,33^ and cytoplasmic drug delivery by polymeric nanomaterials ^34^. Toribio et al. demonstrated that split EGFP-luciferase fusion proteins could be used to measure the cellular uptake of EVs ^35^. However, their assay could not distinguish the cellular uptake of EVs from functional cargo delivery. To our knowledge, this is the first study demonstrating a feasible real-time assay for cytoplasmic cargo delivery by EVs.

Compared to the previously reported assays, the EVCD assay is currently the only method to directly measure the cargo delivery by EVs in living cells. Moreover, the EVCD assay reflected the membrane fusion activity of VSV-G (Fig. 2) and the effect of entry inhibitors (Fig. 3). These results proved the accuracy and feasibility of the assay for quantitative assessment of EV cargo delivery. However, other EV cargo delivery assays may have advantages over the EVCD assay in terms of sensitivity and resolution. For example, RNA (guide RNA or gRNA) delivery by EVs can be measured by a reporter assay utilizing CRISPR/Cas9, the so-called CROSS-FIRE system ^10^. The CROSS-FIRE system can measure the delivery of functional cargo gRNA by EVs at the single-cell level using flow cytometry. This assay is highly sensitive to functional cargo delivery since only a single gRNA delivered to the cytoplasm can lead to the functional readout from recipient cells. However, the CROSS-FIRE system requires multiple additions of EVs to recipient cells and several days are required to obtain the functional readout. Another example of the cargo delivery assay is the use of a BlaM protein as a cargo ^17,24^. Cre recombinase-mediated reporter assay has also been reported in several studies ^11,12,31^. Each assay has its own pros and cons; therefore, comprehensive analysis of EV cargo delivery may expedite the understanding of the mechanism and physiological relevance of EV-mediated cargo delivery.

We estimated that approximately 0.1 fmol of HiBiT per 10^5^ cells, equivalent to approximately 600 molecules of HiBiT per cell, can be detected by the EVCD assay (Fig 2A). EPN-01 proteins spontaneously form a 60-subunit nanocage ^17^; hence, a single nanocage has 60-HiBiT molecules. As described previously, one EV contains 14 EPN-01 nanocages on average ^17^. This indicates that a single EV potentially contains 840 HiBiT molecules (14 × 60 = 840) on average. Together with the estimated sensitivity of the EVCD assay (600-HiBiT/cell), we assumed that only a single event of EV cargo delivery per cell was enough to exceed the detection threshold in the EVCD assay. In spite of the high sensitivity of the EVCD assay, we could not observe cargo delivery of EVs without co-expressing VSV-G (Fig. 2B). This result suggests that the authentic EVs that do not possess known fusion proteins is not capable of delivering the cargo, at least for the combination of HEK293T-derived EVs in recipient HEK293T. It is debatable whether EV-mediated cargo delivery is more efficient in other combinations of EVs and recipient cells.

Fluorescence imaging is usually used to investigate intracellular trafficking of EVs. However, conventional fluorescence imaging of intracellular EVs labeled with fluorescence dyes or fluorescence proteins cannot be used to evaluate cytoplasmic cargo delivery. To overcome the current limitation of fluorescence imaging of EVs, Joshi et al. succeeded in tracing cargo release using fluorescence imaging of the recruitment of fluorescence-labeled galectin or cargo-specific nanobody ^7^. Although their comprehensive analysis is informative to decipher the cargo release process of EVs in recipient cells, it is difficult to distinguish the *bona fide* cargo release from artifacts of galectin recruitment on endosome/lysosomes. Moreover, fluorescence imaging is not feasible for a high-throughput and real-time analysis. Luminescence imaging is more compatible with live cell imaging by avoiding phototoxicity and photobleaching, which are typical issues in live cell imaging. In this study, we succeeded in live cell imaging of an EV cargo delivery (Fig. 5). As discussed above, EPN-01 could form a 60-subunit nanocage, and clustering of HiBiT in nanocage resulted in superior brightness in the imaging, as demonstrated by GFP clustering in a similar protein nanocage ^36^.

Taken together, we developed a quantitative cargo delivery assay of EVs, named the EVCD assay. This assay enabled us to assess the cargo delivery of EVs in recipient cells in real-time. Since EVs are thought to be involved in many biological processes, such as intercellular communication between cells, a feasible EVCD assay may provide insight into the physiological relevance of EVs.

## Methods

### Materials

Drugs and antibodies used in this study are summarized in Supplementary Table 1. NanoLuc substrates were purchased from Promega. The HiBiT peptide (amino acid sequence: VSGWRLFKKIS) and the DrkBiT peptide (amino acid sequence: VSGWALFKKIS) ^22^ were synthesized by GL Biochem.

Additionally, the plasmids used are listed in Supplementary Table 2 and will soon be deposited to Addgene. Plasmids were constructed using PCR-based methods (Gibson Assembly ^37^) and were confirmed by Sanger sequencing.

### Cell culture and transfection

Human embryonic kidney-derived HEK293T (RIKEN Cell Bank) cells were maintained in DMEM supplemented with 10% (v/v) fetal bovine serum (FBS) and 10 μg/mL penicillin-streptomycin. Cells were cultured at 37°C under 5% CO_2_ in a humidified incubator.

One day before the transfection, approximately 2.0 × 10^5^ cells/mL of HEK293T cells were seeded in cell culture dishes or multi-well plates. The following day, HEK293T cells were transfected with plasmid DNA using transfection reagent polyethyleneimine, Transporter 5 Transfection Reagent (Polyscience, Inc.), or branched 25-kDa polyethyleneimine (PEI, Sigma). The ratio of transfection reagent to plasmid DNA was 4:1 (weight). After incubation for 20 to 72 h, cells were subjected to the following experiments.

### Characterization of HiBiT-fused proteins

The expression of HiBiT-fused proteins was analyzed using the HiBiT quantification assay and western blotting. For the quantification of HiBiT-tagged proteins, HEK293T cells transfected with the HiBiT protein expression plasmid were lysed, and the amount of the HiBiT protein was measured using the Nano Glo HiBiT Lytic Detection System (Promega). As a quantification standard for HiBiT proteins, HiBiT peptides was used. For western blotting, HEK293T cells expressing HiBiT-fused proteins were lysed with RIPA buffer containing protease inhibitor cocktail (Nacalai Tesque) and separated by SDS-PAGE. After the blotting of proteins on nitrocellulose membranes, HiBiT-fused proteins were visualized using the Nano-Glo HiBiT Blotting System (Promega). As an internal control of HiBiT proteins, GAPDH in the cell lysate was detected using a conventional western blotting protocol using the same membrane as that used for HiBiT detection.

### PEG precipitation of EVs containing the HiBiT-tagged protein cargo

After 48 to 96 h of transfection, the supernatant from HEK293T expressing cargo HiBiT proteins was collected, centrifuged at 1,500 × *g* for 5 min, mixed with one-third volume of 4 × polyethylene glycol solution (40 w/v%-PEG6000, 1.2 M-NaCl, 1×PBS [pH 7.4]), and incubated at 4°C overnight. The following day, the supernatant was centrifuged at 1,600 × *g* for 60 min, and the residual pellet was resuspended in PBS. Typically, approximately 5 to 10 mL of the supernatant was concentrated to 100 to 200 μL of PBS (approximately 50-fold concentration). The amount of HiBiT proteins in the concentrated EV fraction was measured using the Nano Glo HiBiT Lytic Detection System (Promega). EV marker proteins in the concentrated EV fraction were detected by western blotting as described above.

### Live cell extracellular vesicle cargo delivery (EVCD) assay

Before 24 to 48 h of performing the assay, HEK293T cells (1.0 to 2.0 × 10^4^ cells/well) seeded in a PEI-coated 96-well white plate were transfected with the LgBiT-expressing plasmid. For the EVCD assay, the culture medium of LgBiT-expressing HEK293T cells was replaced with HBSS (+) buffer containing 1 μM DrkBiT, a peptide that complements LgBiT and inactivates LgBiT to reduce the background luminescence signal ^22^. After the addition of the NanoLuc substrate Nano-Glo Live Cell Assay System (Promega) to the cells, a PEG-concentrated EV fraction (approximately 20 to 100 fmol-total HiBiT/well) was added to the cells and monitored for up to 90 min. For the evaluation of inhibitors in the EVCD assay, recipient cells were pretreated with the compounds 1 h before the assay and treatment with the drugs was continued throughout the assay. Microplate of recipient cells was incubated at 37°C and luminescence signal from cells was continually measured by using the Synergy 2 (BioTek) plate reader.

### Live cell luminescence imaging of HiBiT cargo delivery by EVs

For luminescence imaging, HEK293T cells (approximately 1.0 × 10^4^ cells/well) were seeded in a poly-L-lysine (PLL)-coated 35-mm multi-well dish (Matsunami Glass Ind., Ltd.). The following day, cells were transfected with the LgBiT-expressing plasmid and cultured for 24 to 48 h. For live cell imaging, transfected HEK293T cells were washed with HBSS(+) twice and stored in HBSS (+) buffer containing 1 μM DrkBiT and the NanoLuc substrate, followed by the addition of PEG-concentrated EVs (approximately 35 fmol-total HiBiT/well). Continuous live cell imaging was carried out using the software MetaMorph and luminescence microscope LV200 (Olympus) equipped with a 100× objective (Olympus, UPlanSApo, NA = 1.4), a 0.5× relay lens, and an EM-CCD camera, at 37°C. The exposure time for each capture was set at 60 s.

### Statistical analysis

The data in this work were analyzed using one-way ANOVA and post hoc Dunnett’s test or the Student’s *t*-test. All statistical analyses were performed using the Real Statistics Resource Pack software developed by Charles Zaiontz.

## Supplementary Files

- Supplementary Data: Supplementary Methods, Fig. S1, and S2
  Supplementary Methods
  Fig. S1: Immunoprecipitation of EVs containing HiBiT-tagged proteins
  Fig. S2: Requirement of DrkBiT peptide in EVCD assay
- Supplementary Tables: Table S1 and S2
  Table S1: Materials used in this study
  Table S2: Plasmids used in this study
- Supplementary Video
  Time-lapse luminescence imaging of recipient HEK293T cells treated with EVs incorporating VSV-G and HiBiT-tagged EPN-01

## Acknowledgements

We extend our gratitude to Yumi Yukawa for technical assistance in plasmid construction. Experiments using LV200 were supported by Profs. Takeharu Nagai and Mitsuru Hattori at ISIR, Osaka University. All illustrations in this work were created using BioRender.com.

This work was supported in part by JSPS KAKENHI (Grant-in-Aid for Early-Career Scientists 18K18386 and 20K15790 to MS), Research Grant from JGC-Scholarship (to MS), and the “Dynamic Alliance for Open Innovation Bridging Human, Environment and Materials” (MEXT).

