## Supplementary Data for "Real-time luminescence assay for cytoplasmic cargo delivery of extracellular vesicles"

**Supplementary Methods**

*Immunoprecipitation*

Supernatant from transfected HEK293T cells was collected and centrifuged at 1500 ×g for 5 min to remove cell debris. Protein G Mag Sepharose (Cytiva) was washed with PBS, mixed with antibodies (15 µL-beads + 1 µg-antibodies per sample), and incubated for 15 min at RT. Mouse total IgG was used as a negative control of antibodies. After the removal of excess antibodies, supernatant was mixed with antibody-immobilized magnet beads and incubated for 20 min at RT. After the wash of beads with PBS at least 4 times, beads were directly mixed with Nano Glo HiBiT Lytic Detection System (Promega) and the amount of HiBiT was estimated by measuring complemented NanoLuc-derived luminescence using microplate reader Synergy 2 (BioTek).

**Supplementary Figures**

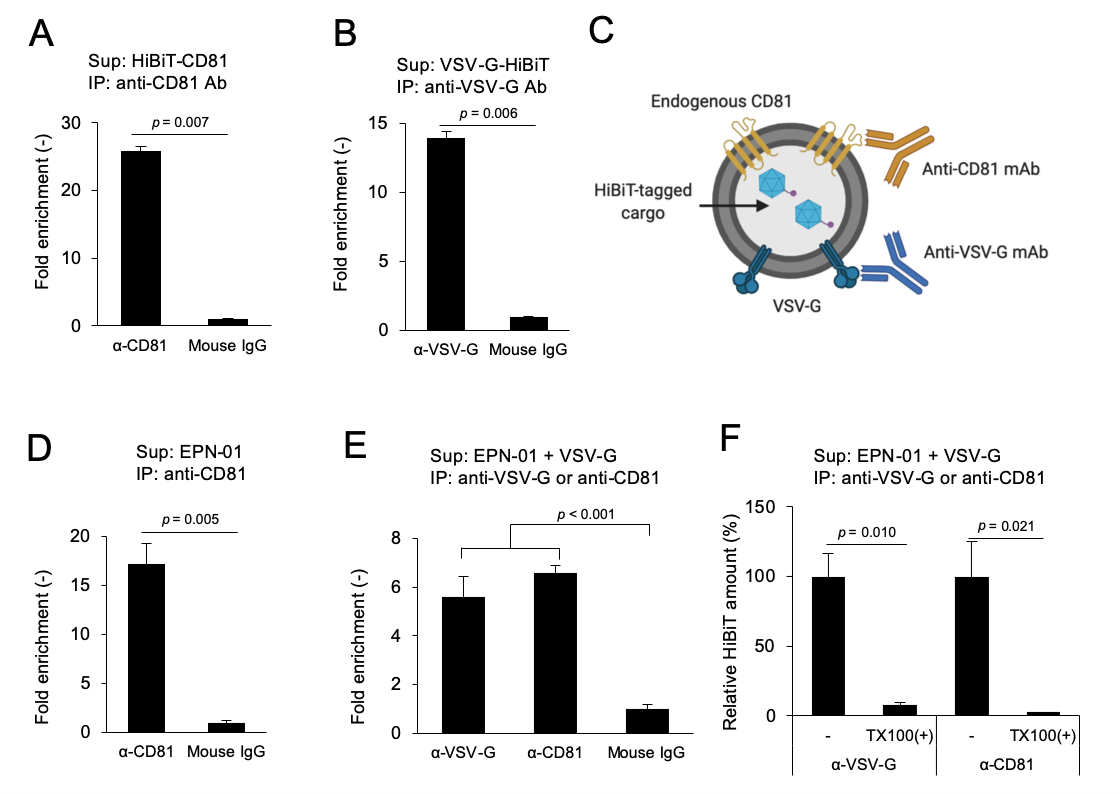

Fig. S1 Immunoprecipitation of EVs containing HiBiT-tagged proteins.

(A) and (B) Validation of anti-CD81 and anti-VSV-G antibody for immunoprecipitation. (C) Scheme of immunoprecipitation of EPN-01-containing EVs. (D) and (E) Immunoprecipitation of EVs containing EPN-01 with or without co-expression of VSV-G. (F) immunoprecipitation of TX100-treated supernatant. Precipitated HiBiT-tagged protein was measured by mixing with LgBiT and NanoLuc substrate. N=3, means ± SD. Student’s t-test for (A), (B), (D), and (F). One-way ANOVA followed by *post hoc* Dunnett’s test for (E).

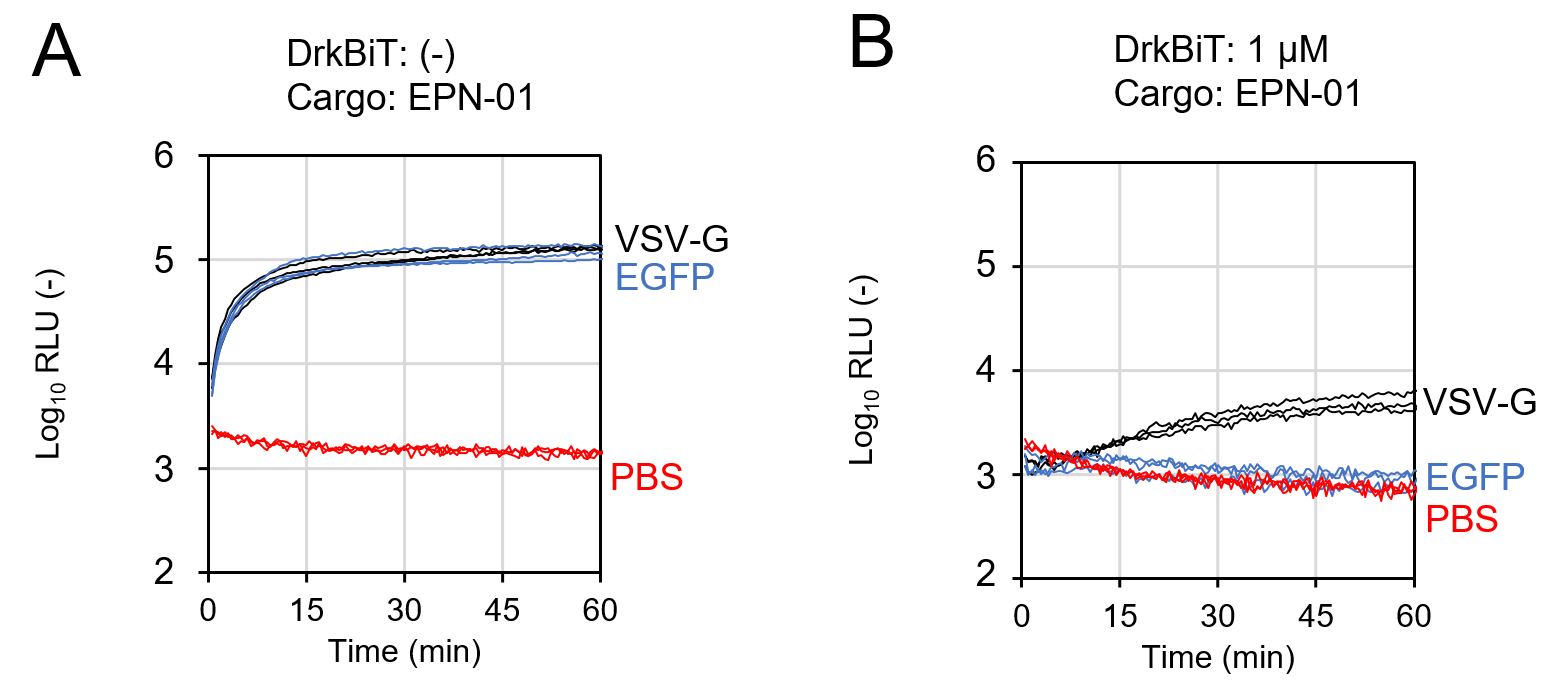

Fig. S2 Requirement of DrkBiT peptide in EVCD assay.

EVs containing HiBiT-tagged EPN-01 (co-expression of either VSV-G or EGFP) were added to LgBiT-expressing recipient HEK293T cells with (A) 0 µM or (B) 1 µM of DrkBiT peptide. PBS was used as a negative control. All kinetics data represent information obtained from experiments conducted in triplicate.
